# Multi-model order spatially constrained ICA reveals highly replicable group differences and consistent predictive results from fMRI data

**DOI:** 10.1101/2022.11.02.514809

**Authors:** Xing Meng, Armin Iraji, Zening Fu, Peter Kochunov, Aysenil Belger, Judy M. Ford, Sara McEwen, Daniel H. Mathalon, Bryon A. Mueller, Godfrey Pearlson, Steven G. Potkin, Adrian Preda, Jessica Turner, Theo G.M. van Erp, Jing Sui, Vince D. Calhoun

**Affiliations:** Tri-Institutional Center for Translational Research in Neuroimaging and Data Science (TReNDS), Georgia State, Georgia Tech, Emory University, Atlanta, GA, USA; Brainnetome Center and National Laboratory of Pattern Recognition, Institute of Automation, Chinese Academy of Sciences, Beijing, China; University of Chinese Academy of Sciences, Beijing, China; Maryland Psychiatric Research Center, Department of Psychiatry, School of Medicine, University of Maryland, Baltimore, MD, USA; Department of Psychiatry, University of North Carolina, Chapel Hill, NC, USA; Department of Psychiatry, University of California San Francisco, San Francisco, CA, USA; San Francisco VA Medical Center, San Francisco, CA, USA; Department of Psychiatry and Biobehavioral Sciences, University of California Los Angeles, Los Angeles, CA, USA; Department of Psychiatry, University of Minnesota, Minneapolis, MN, USA; Departments of Psychiatry and Neuroscience, Yale University, School of Medicine, New Haven, CT, USA; Department of Psychiatry and Human Behavior, University of California Irvine, Irvine, CA, USA; Department of Psychology, Georgia State University, Atlanta, GA, USA; Clinical Translational Neuroscience Laboratory, Department of Psychiatry and Human Behavior, University of California Irvine, Irvine, CA, USA

**Author notes:** **Correspondence Address** Vince D. Calhoun, Tri-Institutional Center for Translational Research in Neuroimaging and Data Science (TReNDS) Georgia State University, 55 Park Pl NE, Atlanta, GA 30303, USA.

**Keywords:** functional network connectivity(FNC), component number, spatially constrained ICA, resting fMRI, machine learning, intrinsic connectivity networks

## Abstract

Brain functional networks identified from resting fMRI data have the potential to reveal biomarkers for brain disorders, but studies of complex mental illnesses such as schizophrenia (SZ) often yield mixed results across replication studies. This is likely due in part to the complexity of the disorder, the short data acquisition time, and the limited ability of the approaches for brain imaging data mining. Therefore, the use of analytic approaches which can both capture individual variability while offering comparability across analyses is highly preferred. Fully blind data-driven approaches such as independent component analysis (ICA) are hard to compare across studies, and approaches that use fixed atlas-based regions can have limited sensitivity to individual sensitivity. By contrast, spatially constrained ICA (scICA) provides a hybrid, fully automated solution that can incorporate spatial network priors while also adapting to new subjects. However, scICA has thus far only been used with a single spatial scale. In this work, we present an approach using scICA to extract subject-specific intrinsic connectivity networks (ICNs) from fMRI data at multiple spatial scales (ICA model orders), which also enables us to study interactions across spatial scales. We evaluate this approach using a large N (N>1,600) study of schizophrenia divided into separate validation and replication sets. A multi-scale ICN template was estimated and labeled, then used as input into spatially constrained ICA which was computed on an individual subject level. We then performed a subsequent analysis of multiscale functional network connectivity (msFNC) to evaluate the patient data, including group differences and classification. Results showed highly consistent group differences in msFNC in regions including cerebellum, thalamus, and motor/auditory networks. Importantly, multiple msFNC pairs linking different spatial scales were implicated. We also used the msFNC features as input to a classification model in cross-validated hold-out data and also in an independent test data. Visualization of predictive features was performed by evaluating their feature weights. Finally, we evaluated the relationship of the identified patterns to positive symptoms and found consistent results across datasets. The results verified the robustness of our framework in evaluating brain functional connectivity of schizophrenia at multiple spatial scales, implicated consistent and replicable brain networks, and highlighted a promising approach for leveraging resting fMRI data for brain biomarker development.

## 1. Introduction

Functional magnetic resonance imaging (fMRI) is a widely used neuroimaging technique to study functional connectivity of the brain. Independent component analysis (ICA) as a widely used approach for assessing brain function can identify maximally independent spatial sources from observed multivariate brain imaging data (Vince D. Calhoun & Adali 2012). The ICA method is able to extract multiple intrinsic connectivity networks (ICNs), i.e., the fMRI independent components (ICs) resulting from group ICA. This ability makes ICA an increasingly attractive data-driven tool to study functional brain networks either at rest or during a cognitive task, including as input into deep learning models (Yan *et al*. 2019).

Group ICA (Vince D. Calhoun *et al*. 2001a, b, Erhardt *et al*. 2011) has been widely applied for investigating functional network biomarkers in neuroimaging data. Group ICA is typically used to extract brain functional networks from multiple subjects in a fully data-driven manner. However, comparison of these networks across datasets can be challenging if they are analyzed separately because the components will be randomly sorted at each run. While one can use regression approaches to estimate the networks in new subjects, this can decrease efficiency as regression neither ensure maximal independence within each subject, nor does it fully adapt the spatial networks to the new data (Erhardt *et al*. 2011). Consequently, when leveraging high er-order statistics, previous work has shown the statistical and classification performance of regression-based approaches are reduced compared to the ICA solutions (Salman *et al*. 2019a).

In contrast, spatially constrained ICA (scICA) of fMRI (Lin *et al*. 2010, Du & Fan 2013) were proposed to overcome difficulties in identifying components of interest and determining the optimal number of components in ICA analysis because scICA incorporates available spatial prior information about the sources into standard blind ICA. Previous studies (Sui *et al*. 2009, Qi *et al*. 2018, Salman *et al*. 2019b) have demonstrated the benefit of using scICA in functional MRI analysis. Recent work has proposed deriving component priors by identifying replicable component maps across multiple large datasets analyzed separately, then using these templates in a spatially constrained ICA. This template, called ‘Neuromark’, has been used in a variety of studies to date (Du *et al*. 2019, 2020), which has multiple strengths: 1) fully automated, avoiding the need to select and label components, 2) adapts the component maps and timecourses to individual subjects, making cross-validation easier, and 3) provides comparability across studies. However, scICA has to date only been applied with priors at a single spatial scale (i.e., model order). In previous work we have shown that the neuromark approach (using the multi-objective optimization ICA with reference algorithm, or MOO-ICAR) estimates subject-specific ICs that provide more optimal independence and better spatial correspondence across different subjects and achieve higher spatial and temporal accuracy compared to the existing ICA methods. Multiple studies have used this approach today. For example, we have used the Neuromark IC template to estimate subject-specific networks that were then used as features in a support vector machine (SVM)-based framework to predict response to medication (either antidepressants or mood stabilizers) in bipolar disorder and major depressive disorder patients (Osuch *et al*. 2018, Du *et al*. 2019, 2020).

However, such work focuses on priors derived from a single ICA model order (i.e., 53 networks from 100 components estimated), ignoring the importance of capturing functional information at different levels of spatial granularity as well as the between-order information. In recent work, we have shown the advantage of working with multiple spatial scales (i.e., multi model-order ICA) (Iraji *et al*. 2021, Meng *et al*. 2021) Multi-model order ICA provides a comprehensive way to study brain functional network connectivity within and between multiple spatial scales, highlighting findings that would have been ignored in a single model order analysis. Previous studies have highlighted the benefit of studying functional network connectivity within and between multiple spatial scales(IC orders) in schizophrenia (A. Iraji et al. 2021, Meng, Xing et al. 2021).

On the other hand, schizophrenia is a severe psychiatric disease characterized by hallucinations, delusions, loss of initiative, and cognitive dysfunction (Van Den Heuvel & Fornito 2014). Schizophrenia is a heterogeneous syndromic diagnosis of exclusion, lacks unique symptoms, and is diagnosed clinically by both positive and negative symptoms (American Psychiatric Association (2013)). Schizophrenia has been hypothesized as a developmental disorder of disrupted brain function, which can be characterized by functional dysconnectivity and changes in functional integration (Friston & Frith 1995, Stephan *et al*. 2006). Therefore, studying functional network connectivity can provide important information about brain functional integration and its schizophrenia changes, potentially improving our understanding of the actual brain pathology underlying different schizophrenia subcategories (A. Iraji et al. 2021). However, most previous research has focused on studying brain functional connectivity of schizophrenia within a single model order, ignoring the importance of cross-connectome information at different spatial scales. Thus, using ICNs resulted from spatial constraint ICA with multiple model orders can fill the gap.

To the best of our knowledge, no studies have shown the robustness of spatially constrained ICA in the context of capturing group differences at single or multiple spatial scales. In this study, we proposed a framework to evaluate multi-model order scICA using MOO-ICAR to capture robust, replicable schizophrenia-related alterations and identify potential biomarkers for schizophrenia classification. We used a recently developed multi-model order ICA template (Meng, Xing *et al*. 2021) and together with the proposed scICA based framework to identify consistent predictive ICNs in schizophrenia. To evaluate the robustness of our framework, we built our framework on one dataset and evaluated the performance on an independent dataset.

## 2. Methods

### 2.1 Dataset and preprocessing

We used two datasets in this study. The first data set, ‘dataset 1’, was used as a discovery and validation dataset, and the second ‘dataset 2’ was used as a replication dataset. Dataset 1 was mainly used to extract predictive features based on which classification was built and validated. Dataset 2 was used to train the classification model using the features that were selected from dataset 1.

Dataset 1 was selected from three different studies, one with seven sites (fBIRN: Functional Imaging Biomedical Informatics Research Network), one with three sites (MPRC: Maryland Psychiatric Research Center), and one single site (COBRE: Center for Biomedical Research Excellence). This resulted in a total 827 individuals, including 477 subjects (age: 38.76 ± 13.39, females: 213, males: 264) of typical controls (TC) and 350 schizophrenia individuals (age: 38.70 ± 13.14, females: 96, males: 254). The parameter settings for the resting-state fMRI (rsfMRI) data collected in the fBIRN data were the same across all sites, with a standard gradient echo-planar imaging (EPI) sequence (repetition time (TR)/echo time (TE) = 2000/30 ms, voxel spacing size = 3.4375 × 3.4375 × 4 mm, field of view (FOV) = 220 × 220 mm, and a total of 162 volume). Six of the seven sites used 3-Tesla Siemens Tim Trio scanners, and one site used a 3.0 Tesla General Electric Discovery MR750 scanner. For COBRE data, rsfMRI images were acquired using a standard EPI sequence (TR/TE = 2000/29 ms, voxel spacing size = 3.75 × 3.75 × 4.5 mm, FOV = 240 × 240 mm, and a total of 149 volumes. Data were collected using a 3-Tesla Siemens Tim Trio scanner. The MPRC dataset were acquired using a standard EPI sequence in three sites, including Siemens 3.0 Tesla Siemens Allegra scanner (TR/TE = 2000/27 ms, voxel spacing size = 3.44 × 3.44 × 4 mm, FOV = 220 × 220 mm, and 150 volumes), 3.0 Tesla Siemens Trio scanner (TR/TE = 2210/30 ms, voxel spacing size = 3.44 × 3.44 × 4 mm, FOV = 220 × 220 mm, and 140 volumes), and 3.0 Tesla Siemens Tim Trio scanner (TR/TE = 2000/30 ms, voxel spacing size = 1.72 × 1.72 × 4 mm, FOV = 220 × 220 mm, and 444 volumes). This data has also been used in prior work (A. Iraji *et al*. 2021, Meng, Xing *et al*. 2021).

Dataset 2 contained a total of 815 subjects, collected from several Chinese hospitals, including 326 subjects (age: 29.81 ± 8.68, females: 167, males: 159) of typical controls and 489 SZ individuals (age: 28.98 ± 7.63, females: 229, males: 260). The subjects were Chinese ethnic Han groups. The dataset was recruited from seven sites in China with the same recruitment criterion, including Peking University Sixth Hospital; Beijing Huilongguan Hospital; Xinxiang Hospital Simens; Xinxiang HospitalGE; Xijing Hospital; Renmin Hospital of Wuhan University; Zhumadian Psychiatric Hospital (Yan *et al*. 2019). The resting-state fMRI data were collected with the following three different types of scanners across the seven sites: 3.0 Tesla Siemens Tim Trio Scanner, 3.0 T Siemens Verio Scanner, and 3.0 T Signa HDx GE Scanner (TR/TE = 2000/30 ms, voxel spacing size = 3 × 3 × 3 mm, FOV = 220 × 220 mm, and 480/360 volumes). Subjects were instructed to relax and lie still in the scanner while remaining calm and awake.

The two datasets were preprocessed according to the same procedures as in our previous study (A. Iraji et al. 2021). To summarize, preprocessing was mainly performed using the statistical parametric mapping (SPM12, https://www.fil.ion.ucl.ac.uk/spm/) toolbox. First, we discarded the first five volumes for magnetization equilibrium. We then performed rigid body motion correction using the toolbox in SPM to correct subject head motion, followed by the slice-timing correction to account for timing difference in slice acquisition. For each subject, the translation of head motion was less than 3 mm and the rotation of head motion was less than 3° in all axes through the whole scanning process. And the next step, the rsfMRI data of each subject was subsequently warped into standard Montreal Neurological Institute (MNI) space using an echo-planar imaging (EPI) template and smoothed using a Gaussian kernel with a 6 mm full width at half-maximum (FWHM = 6 mm). The voxel time courses were z-scored (variance normalized). To make it consistent, the minimum data length (135 volumes) across all subjects from the two datasets was selected for further analysis.

### 2.2 Analysis framework

The framework to explore group differences and identify predictability of PANSS scores (Meng, Xing *et al*. 2021) in schizophrenia is provided in Fig.1. There are three major components in this framework: 1) apply spatially constrained ICA to extract corresponding functional regions and time-courses (TCs), calculate msFNC matrix for each individual; 2) perform feature selection based on the msFNC matrix, and build the SVM classification model; 3) identify predictive ICN domains, and evaluate differences between schizophrenia and control groups.

**Figure. 1.**
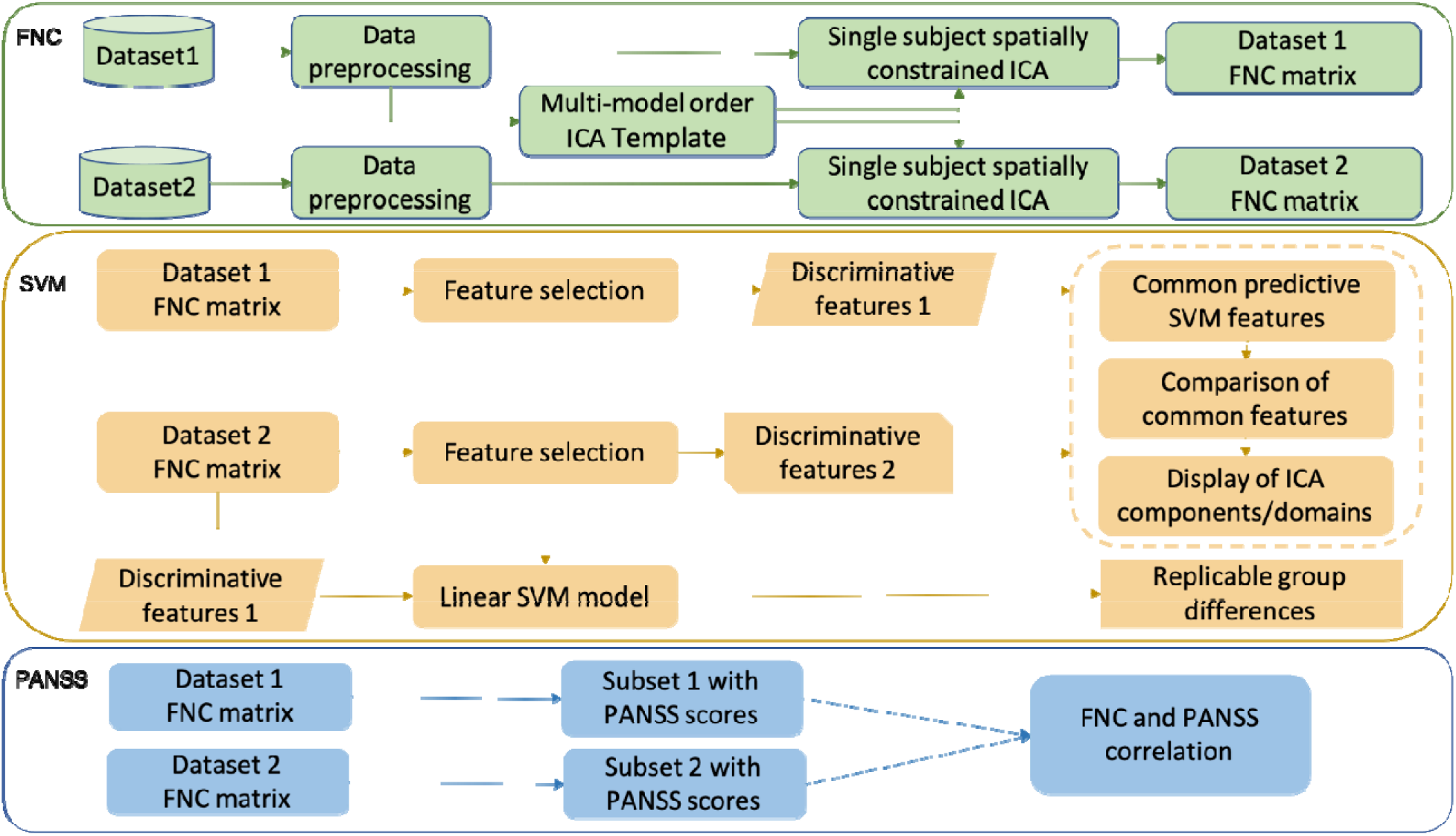
Workflow of our framework. After preprocessing, we calculated the msFNC matrix based on the spatially constrained ICA for both datasets. We then performed feature selection on the msFNC matrix of dataset 1, to find out predictive ICN features. And the next step, we built the SVM model on dataset 2, using the ICN features selected from dataset 1. To find out general predictive ICN features across the different dataset, we repeated the feature selection process on dataset 2, and compared them with the predictive features selected from dataset 1. Thus, we identified consistent predictive features across different datasets. And finally, we extracted a subset of SZ with available symptom scores from dataset 1 and dataset 2, and calculated the linear correlation between msFNC and the symptom score. We then made a comparison between them.

### 2.3 Spatially constrained ICA analysis

The proposed approach is based on scICA, which incorporates a spatial reference in the ICA algorithm. This allows us to extract only desired ICs, and hence we do not need to run a complete ICA to extract all sources. The scICA analysis was performed using the GIFT software (http://trendscenter.org/software/gift, Vince D. Calhoun *et al*. 2001a, Vince D. Calhoun & Adali 2012, Iraji *et al*. 2020, 2021). We ran scICA on each subject for both dataset 1 and dataset 2, using a template that contained a total of 127 ICNs, that were determined from multiple model orders from 25, 50, 75, and 100. This template was obtained and labeled using dataset 1 (A. Iraji *et al*. 2021). We identified 15, 28, 36, and 48 ICNs from each model order of 25, 50, 75, and 100, respectively.

### 2.4 msFNC analysis on spatially constrained ICA

msFNC was computed between each pair of ICN time courses by calculating the Pearson correlation coefficient between ICN timecourses (V D Calhoun *et al*. 2003, Madiha J Jafri *et al*. 2008, Allen *et al*. 2011), which resulted in a 2D symmetric ICN × ICN msFNC matrix for each individual. Each cell of the msFNC matrix represented the functional connectivity between two ICNs. To capture functional interaction across different spatial scales, we calculated the functional network connectivity between each pair of ICNs across all model orders. ICN time courses were interpolated to 2s for a subset of dataset 1 (15%) with a sampling rate other than 2 s and for all of dataset 2. We aggregated the msFNC matrix from all subjects into an augmented 2D matrix. We then calculate the mean msFNC matrix of all subjects for further analysis.

### 2.5 Feature selection on msFNC

The feature selection process was performed using only dataset 1. Each msFNC pair was considered as the input feature for classification, and the category of group TC or SZ was considered as the response vector. Given that some of those msFNC features might be non-informative or redundant for classification, we performed feature selection using Relief (Verma & Salour, Al 1992) to improve classification performance and speed up computation. Relief ranks predictors with *k* nearest neighbors (we set *k* to 10 as the most used value). The function returns the indices of the most important predictors and the weights of the predictors. Feature selection was carried out before classifier training through the recursive feature elimination step. For each round of feature selection, 50% of the training data was selected. We repeated it for ten rounds and retained those features with a high average weight (top 70%) among all the rounds. We thus narrowed the set of features to a subset of the original feature set, eliminating msFNC features’ redundancy. The classification model was built based on the selected msFNC feature set.

### 2.6 Support vector machine-based classification (SVM)

The SVM (Verma & Salour, Al 1992) is a widely used binary classification method due to its ability to deal with high-dimensional data and versatility in modeling diverse sources of data. The SVM has been widely applied in numerous neuroimaging classification studies and has achieved remarkable results due to its excellent generalization performance. Our motivation for using SVM over other approaches was due to its sensitivity, resilience to overfitting, ability to extract and interpret features, and superior performance in fMRI data classification (De Martino *et al*. 2008, Pereira *et al*. 2009, Ecker *et al*. 2010, Liu *et al*. 2013, Vergun *et al*. 2013, M. Wang *et al*. 2019, Saha *et al*. 2021). To investigate the group differences, we built a binary SVM classifier using a linear kernel(Chih-Wei Hsu *et al*. 2003), as a straightforward baseline classifier, to demonstrate the practicability of our framework. We also compared the performance of SVM and random forest as a comparison model on a small training set, as SVM outperformed random forest in that case, we utilized SVM as our baseline model.

We built an SVM model on dataset 2, using the ICN features selected from dataset 1 as mentioned in the previous feature selection section. Thus, to verify the robustness of our model. To obtain stable performance, we iteratively built and evaluated the classification model multiple times on dataset 2. For each iteration, we randomly split the whole dataset 2 into 80% of the training set and 20% of the testing set. The test set was held out for final evaluation. We ran the modeling process for a total number of 100 iterations and evaluated the SVM model based on average specificity, sensitivity, and F1 score (the harmonic mean of the precision and recall) across all iterations.

### 2.7 Compare common predictive features

To identify common predictive ICN features in detecting schizophrenia across different datasets, we repeated the same feature selection procedure on dataset 2, and compared the selected features from dataset 2 with the ones that were selected from dataset 1. As we have a large number of features (in total 8001 msFNC features, between 127 pairs of ICNs), we wanted to focus on those highly predictive features, and explore a heuristic result. As a result, we selected around 1% (96 features out of 8001 in total) of top-ranked features from each of the datasets. We then compared the two feature sets and selected the overlapping features. We normalized their feature weights for further comparison. The major goal in this section is to find out consistent predictive ICN features across different datasets, and thus to show the robustness of our study.

### 2.8 Correlation between msFNC and Symptom score

We investigated the correlation between msFNC and the symptom scores, measured by the positive and negative syndrome scale (PANSS) (Kay *et al*. 1987). We extract 143 schizophrenia individuals from dataset 1, and 149 schizophrenia individual subjects from dataset 2 separately, with valid symptom scores (PANSS total, PANSS positive, PANSS negative). We then calculated the linear correlation between the msFNC and the symptom scores for each of the subjects of the two datasets and made a comparison between them.

## 3. Results

### 3.1 msFNC analysis across different databases

Fig. 2 presents the mean msFNC (z-fisher score) between 127 ICNs of all model orders, for dataset 1 and dataset 2, separately. The overall patterns of the mean msFNC matrix of dataset 1 and dataset 2 were similar. We observed that the cerebellum (CR), somatomotor (SM), subcortical (SB), temporal (TP), and visual (VS) were highly correlated with themselves in msFNC for both of the datasets, which were more homogeneous than the default mode (DM) and cognitive control (CC) domains.

**Fig. 2.**
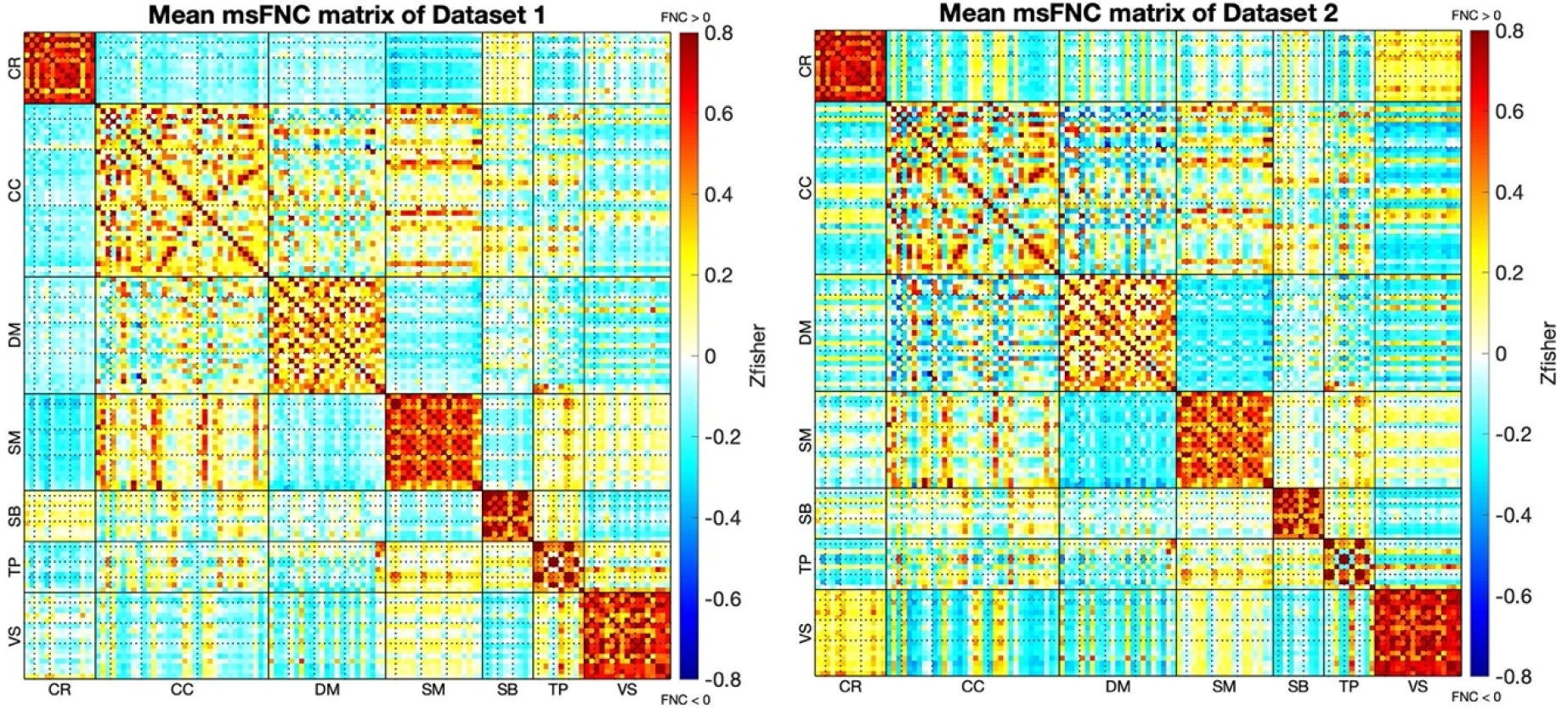
Mean msFNC plot of dataset 1 (left) and dataset 2 (right). We calculated the mean msFNC (z-fisher score) based on the aggregated msFNC matrix of all individuals. The ICNs in these msFNC matrices were sorted by domains first, and within each domain, ICNs were sorted by model orders (from 25 to 100). The dot lines in each domain divide different model orders. ICNs were sorted in the order of cerebellum (CR), cognitive control (CC), default mode (DM), somatomotor (SM), subcortical (SB), temporal (TP), and visual (VS). The overall pattern of the two datasets is similar. Stronger correlations in CR vs. VS and anticorrelations in DM vs. SM, CC vs. DM were seen in dataset 2, and stronger anticorrelations in CR vs. SM in dataset 1.

We then evaluated the group difference between SZ and TC groups for the two datasets. The most dominant increases in msFNC in the SZ group for both datasets were mainly seen between the ICNs of cerebellum vs. somatomotor, cerebellum vs. temporal, and cerebellum vs. visual. Increases in msFNC were also seen between the ICNs of subcortical vs. visual, subcortical vs. temporal, and subcortical vs. somatomotor. In addition, both datasets show a relatively large decrease in the SZ groups compared to the TC groups between the ICNs of the subcortical vs. cerebellum, somatomotor vs. visual, somatomotor vs. temporal. The overall group differences between SZ and TC for the two datasets are shown in Fig 3.

**Fig. 3.**
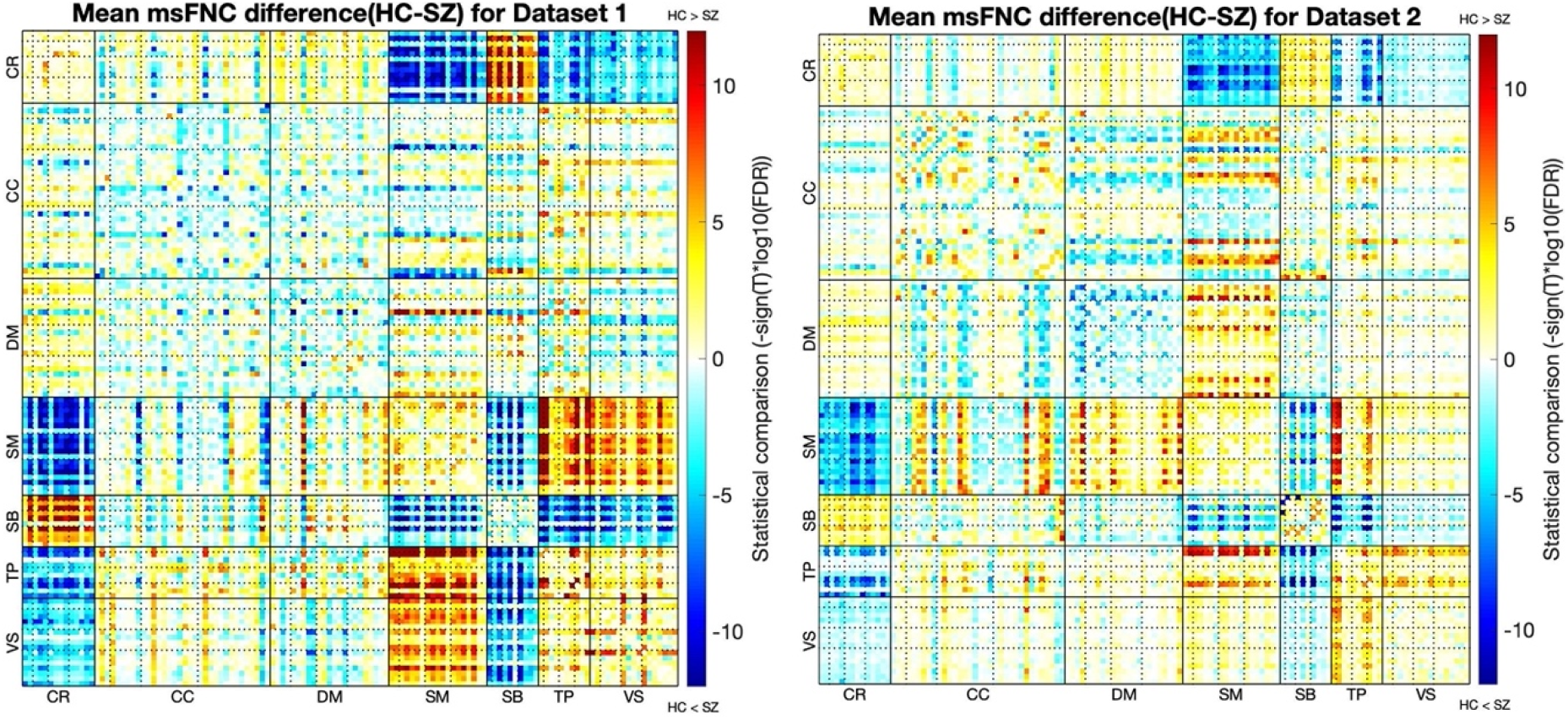
Mean msFNC difference (TC - SZ) matrix for dataset 1 (left) and dataset 2 (right). This figure shows the difference between TC and SZ in msFNC. Blue areas indicate increased msFNC in SZ compared to TC, and red areas indicate decreased msFNC in SZ compared to TC. The intensity values (-sign(T)*log10(FDR)) have been calculated. Increases in msFNC were seen in CR vs. SM, CR vs. VS, CR vs. TP, SB vs. VS, SB vs. TP, SM vs. SB for both datasets. And decreases in msFNC were mainly seen in CR vs. SB, SM vs. VS and SM vs. TP. Discernible group differences were observed in dataset 1, compared to dataset 2, as it shows in the darker regions.

As we mentioned, we selected around 1% of top-ranked features from each of the datasets and then compared the two feature sets and retained the overlapping features, which resulted in 18 common predictive features. Fig. 4 shows the Connectogram of the average msFNC difference between TC and SZ for the 18 common predictive features between the two datasets. The common predictive features have shown distinct group differences between TC and SZ in msFNC for both datasets. The selected common features were considered the most predictive features in detecting group differences in schizophrenia, as they were selected separately from each dataset by feature selection. These common features identified ICN domains that are highly related to schizophrenia. It is observed that the common predictive features also show consistent group differences and consistent signs (positive vs negative) for the msFNC in the two datasets.

**Fig. 4.**
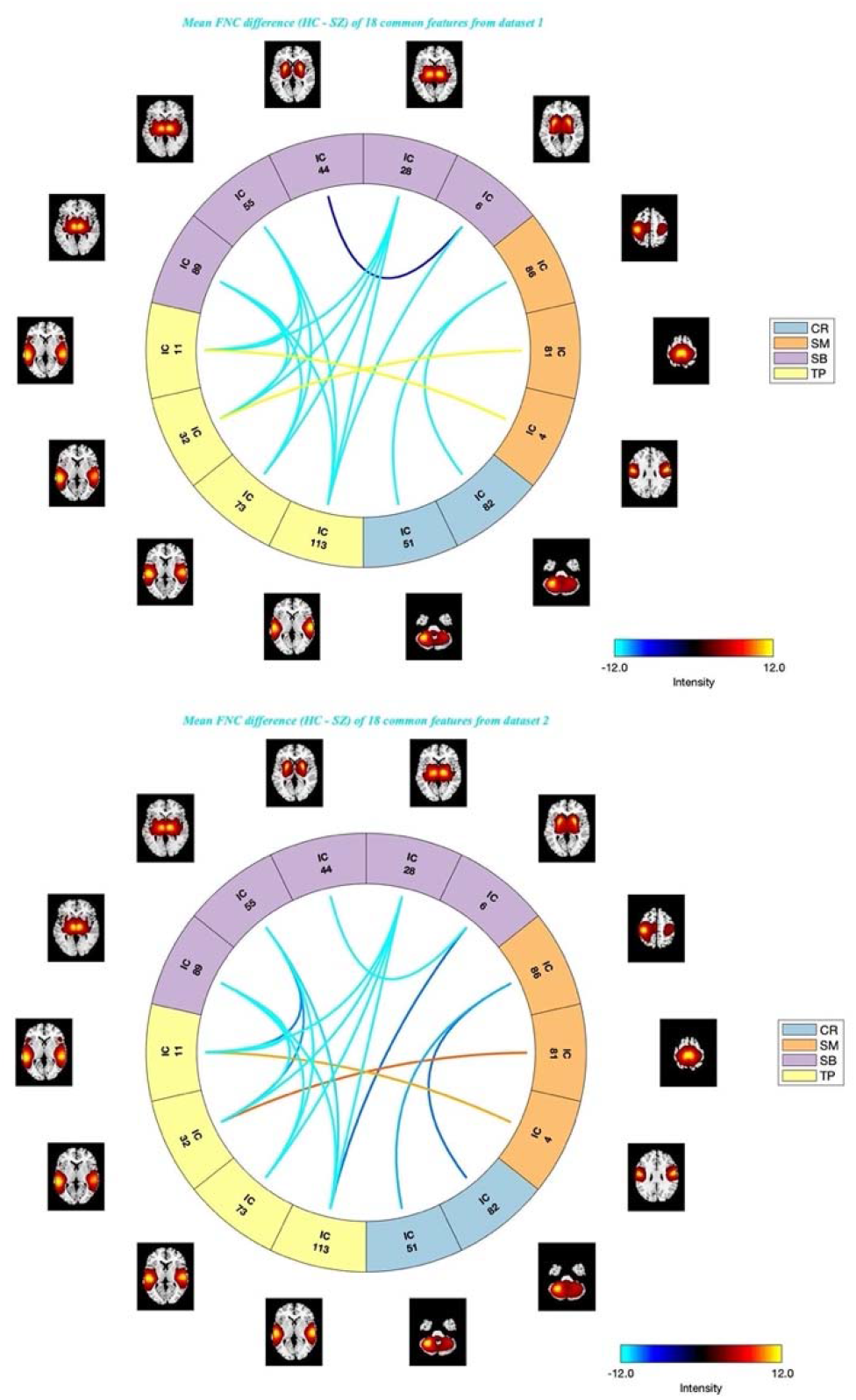
Connectogram of mean msFNC difference (TC - SZ) matrix of 18 common predictive features for dataset 1 (left) and dataset 2 (right). The common features show strong increases in msFNC in the domains of SB vs. TP, SB vs. SB, CR vs. SM, and decreases in SM vs. TP, for both of the datasets.

### 3.2 Support vector machine-based classification (SVM)

We evaluated the performance of the SVM model as shown in Table 1. As we mentioned, the SVM model was built using the predictive features selected from dataset 1, and then trained and verified on dataset 2. The performance was averaged across 100 iterations of modeling. As shown in the table, the average accuracy of the classification model was 81.4%, with a precision of 82.6%, 87.6% recall, and 84.9% F1 score.

**Table 1.**
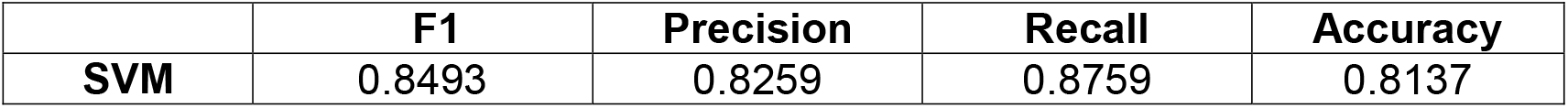
The average performance of the SVM model for 100 iterations.

To better understand the ICN domains that the 18 common predictive features connected and the features’ contribution in detecting group differences, we show the connectograms (as shown in Fig. 5) of the feature weights of the common features. The majority of the common features fell into the ICNs domains of subcortical vs. temporal, the rest of them fell into the ICNs domains of somatomotor vs. temporal, somatomotor vs. cerebellum, and within the subcortical, which may indicate an important role of these domains in detecting schizophrenia. The lines within the circles represent ICN features, and the outer circles indicate the ICN domains the features belong to. As we mentioned, the common features fell into four ICN domains, cerebellum, somatomotor, subcortical, and temporal, with 2, 3, 5, and 4 ICs in each of them separately. It is observed that the majority ICs (11 out of 14) involved were from higher model orders (75∼100), and most of them were from between model orders. The subfigure Fig. 5c the averaged feature weights of the common features of the combined two datasets.

**Fig. 5.**
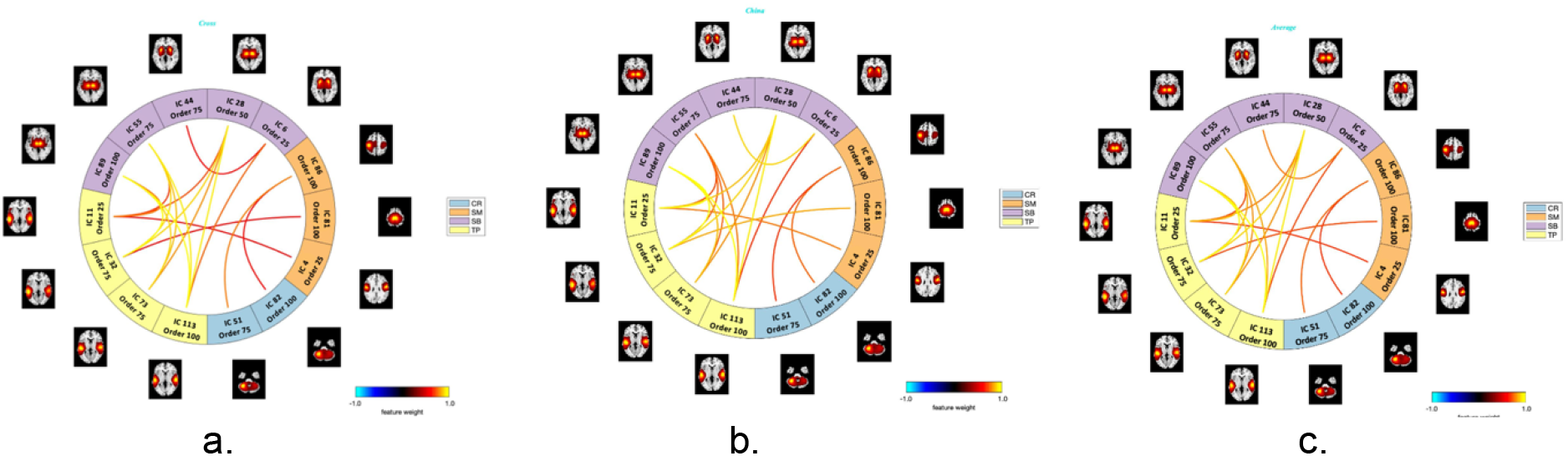
Connectogram of feature weights of the 18 common features, for dataset 1 (a), dataset 2 (b), and the combined dataset (c). The common features have shown strong predictive strength in predicting group differences of TC and SZ, for both of the datasets. Statistically, the relevance level of a relevant feature is expected to be larger than zero and that of an irrelevant one is expected to be zero (or negative). The SB, TP, SM, CR, SM contribute most to the classification, for both of the datasets. The ICN features in domain TP (yellow) vs. SB (purple) generally have higher weights compared to other ICN domains, which indicate their predictive strength in detecting group differences of TC and SZ.

### 3.3 Correlation between msFNC and symptom scores

In order to detect the correlation between msFNC and symptom scores, we calculated the linear correlation between the msFNC of each subject and its symptom scores (including PANSS total, PANSS positive, and PANSS negative). The following figures show the correlation matrix between msFNC and symptom scores in the two datasets. It is observed that the correlation in the ICN domain of somatomotor vs. cerebellum in dataset 2 was noticeably stronger than that of dataset 1. And also, dataset 2 shows generally stronger anticorrelations between msFNC and symptom scores in the ICN domains of somatomotor vs. visual, somatomotor vs. temporal, and somatomotor vs. cerebellum, compared to dataset 1, which have indicated the major differences of the two datasets. In addition, both of the datasets have shown strong anticorrelations within the subcortical domain between the correlation of msFNC and PANSS positive. We then calculated the correlation between msFNC and symptom scores with the combined dataset of the two datasets, as shown on the third row of Fig. 6 (Appendices).

**Fig. 6.**
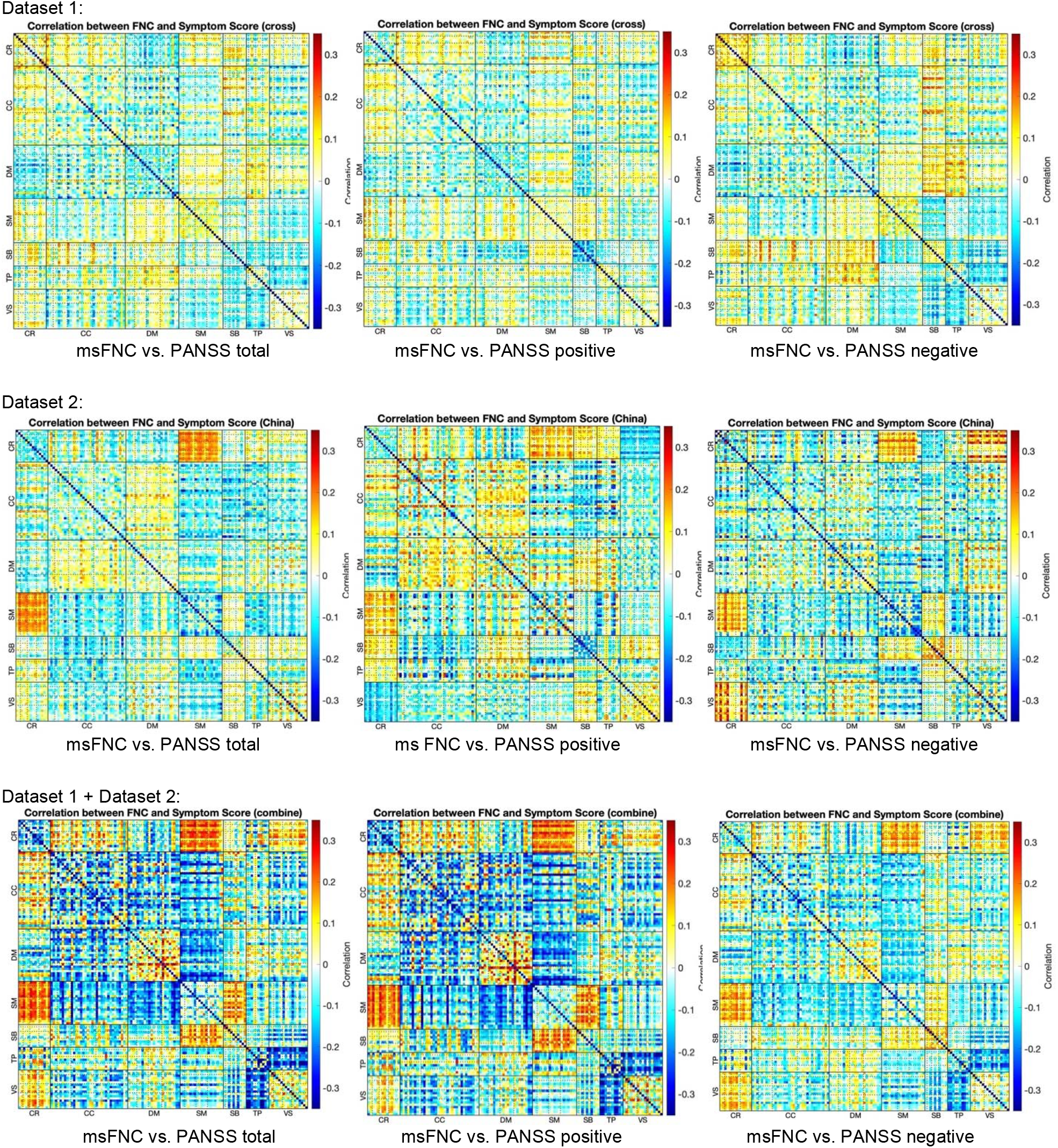
Correlations between msFNC and symptom score, for dataset 1 (the first row), dataset 2 (the second row), and the combined dataset of the two datasets (the third row). Each row shows the correlations between msFNC and the PANSS total (left), PANSS positive (middle) and PANSS negative (right), separately. We extracted 143 and 149 SZ with valid PANSS(total, positive and negative) scores from dataset 1 and dataset 2 separately. We calculated linear correlations between the msFNC matrix and the PANSS scores for each SZ patient. It is noticeable that the correlations between msFNC and the symptom scores in the ICN domain of CR vs. SM are stronger in dataset 2, compared to dataset 1.

Fig. 7 shows the connectograms of the correlation between msFNC and symptom scores for the 18 common predictive features selected from the two datasets. The common predictive features in dataset 2 (the second row) have shown stronger correlations between msFNC and symptom scores (PANSS total on the left, PANSS positive in the middle, and PANSS negative on the right) in the ICN domain of somatomotor vs. cerebellum, compared to the ones in dataset 1 (the first row). The third row of the figure shows the correlation between msFNC and symptom scores of the common features combined of the two common feature sets.

**Fig. 7.**
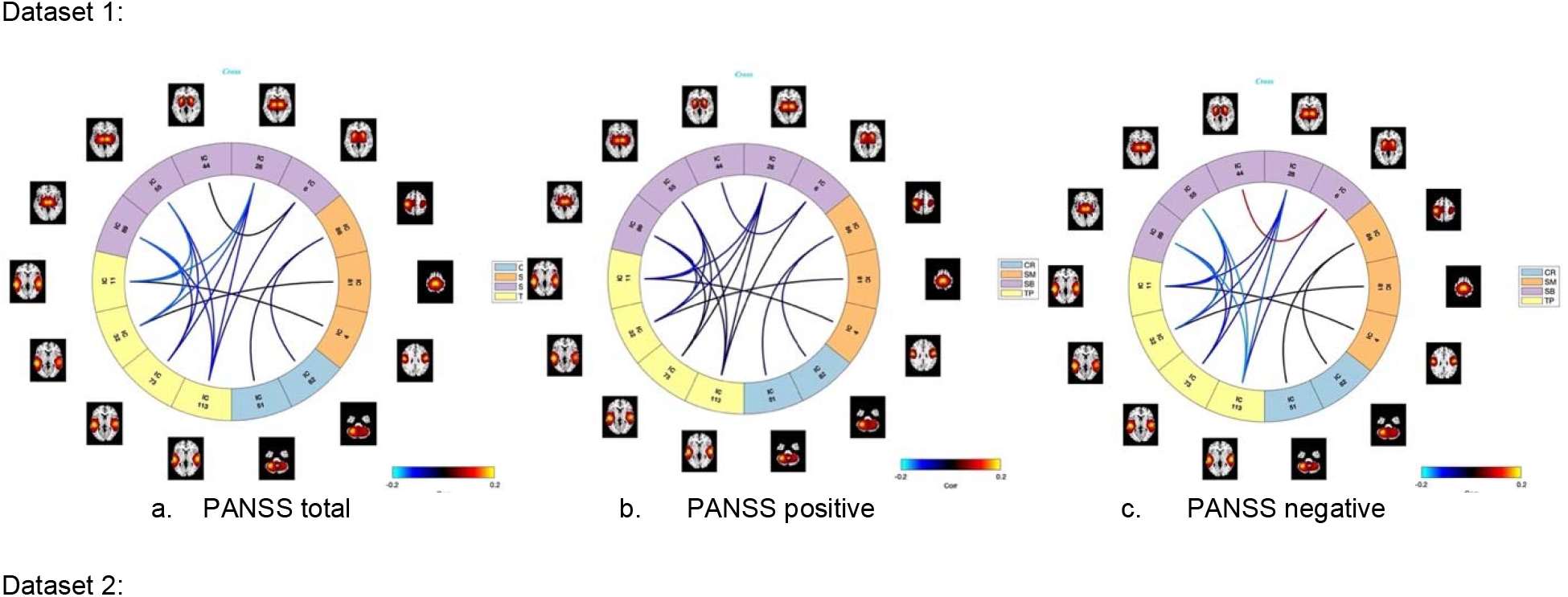
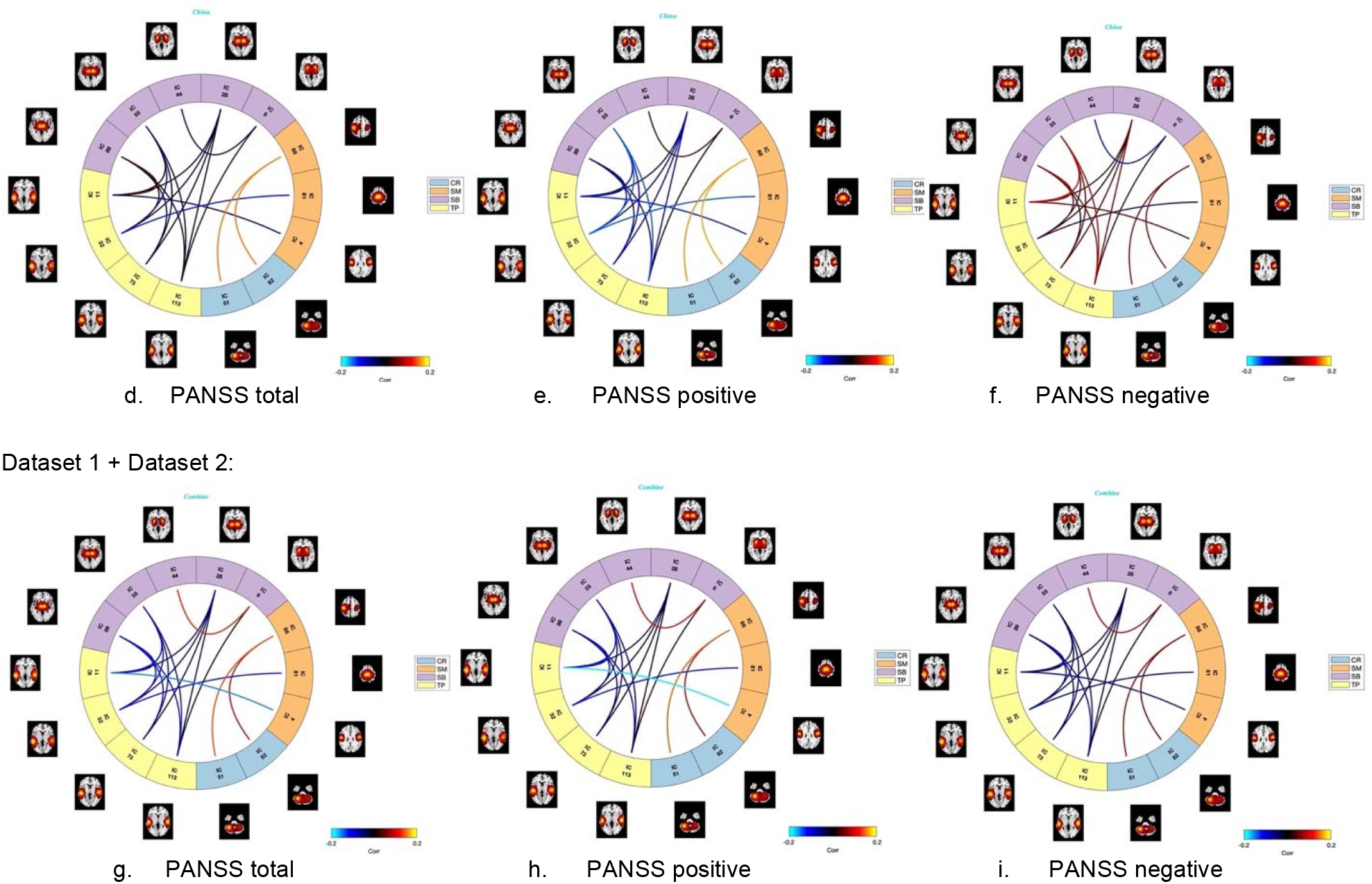
Connectograms of the correlation between msFNC and symptom scores for the 18 common features. The connectograms of the first row and second row show the correlations between msFNC and the PANSS total (a. and d.), PANSS positive (b. and e.) and PANSS negative (c. and f.), for dataset 1 and dataset 2 separately, where PANSS total is the sum of PANSS positive and PANSS negative. The connectograms of the last row are the correlations between msFNC and the PANSS total (g.), PANSS positive (h.) and PANSS negative (i.) for the combined dataset. Linear correlation was calculated between the msFNC matrix and the PANSS scores for each SZ patient. It is seen that the anticorrelations between the ICN domains of TP (yellow) vs. SB (purple) were generally stronger in dataset 1 (the first row) compared to dataset 2 (the second row), as they show in the lighter blue lines within the circles. Stronger correlations between CR (blue) vs. SM (orange) in dataset 2 were also noticeable, as they show in the red lines in the second row, compared to the first row of dataset 1.

## 4. Discussion

In this work, we present a framework to extract subject-specific intrinsic connectivity networks from fMRI data at multi-model order spatially constrained ICA. We built our predictive model based on the ICN features selected from one dataset, and trained the model on a different dataset, using the predictive features selected from the first dataset. We then compared the two independent datasets, regarding their msFNC patterns, predictive ICN features, group differences between typical controls and schizophrenia patients, and also the correlations between msFNC and symptom scores. Results have shown consistent predictive strengths of the four ICN domains of the cerebellum, somatomotor, subcortical, and temporal domains, in detecting schizophrenia. The multiscale ICA template was generated from dataset 1. It was also used as a training set of feature selection for the classification model. Dataset 2 was used as an independent dataset on which the classification model was built and tested. The performance of the classification model reached up to 85% F1 score, 83% precision, and 88% recall. Results suggest the proposed framework would probably provide similar performance if applied on other datasets with different demographics (race, age, gender, etc.), given the replicable evidence we obtained. Our results demonstrated that MOO-ICAR is capable of obtaining subject-specific ICNs with strong independence, which in the meanwhile reduced the computational cost compared to the standard ICA methods.

As we observed, the 18 common predictive features fell into four ICN domains, cerebellum, somatomotor, subcortical, and temporal. And the majority ICs involved were from between model orders, for example, subcortical vs. temporal, somatomotor vs. temporal, somatomotor vs. cerebellum. The results have shown the importance of studying brain functional connectivity at cross-spatial scales.

There are some limitations of our study that we think might be improved in the future. First, for simplicity, we only selected and evaluated a small set (the top 1.2%, which resulted in 96 features out of 8001 in total) of top-ranked common ICN features from the two datasets as our preliminary results. The selected 18 common features have demonstrated their consistent strengths in detecting group differences between TC and SZ groups in schizophrenia. However, it is worth exploring a larger range of scales of the common feature set in future work and evaluating the optimal size of the common predictive feature set to build the classification model. This may improve the performance of the predictive model. Furthermore, we only tried the SVM model as the classification model in this work to provide a baseline, many other more advanced classification models could be used, including deep learning models. We would like to compare other classification models with the one we built in this study in the next step. In this work, we were performing feature selection on the labeled dataset. As future work, we can explore some unsupervised feature selection methods, such as filter methods (Noelia Sánchez-Maroño,*et al*. 2007), wrapper methods (Yang *et al*. 2013) or embedded methods (S. Wang *et al*. 2015), which can be used for unlabeled data, to find the best set of features to build the predictive models.

## 5. Conclusions

We reported a new framework for detecting FNC differences between groups at multiple spatial scales using spatially constrained ICA. The results showed consistent evidence of the four ICN domains cerebellum, somatomotor, subcortical and temporal, especially in detecting aberrant FNCs in schizophrenia on two independent datasets. These results highlight replicable cross-spatial scale msFNC differences which may inform our understanding of the neural patterns linked to schizophrenia. Future work might focus on further replication and potentially focus on interventional approaches targeting the highlighted domains.

## Acknowledgments

Thanks to the studies which collected the original data.

## Authorship Confirmation Statement

Xing Meng: Conceptualization; Formal analysis; Investigation; Methodology; Visualization; Writing – original draft; Writing – review & editing. Armin Iraji: Conceptualization; Investigation; Methodology; Visualization; Writing – review & editing. Zening Fu: Data collection and preprocessing. Peter Kochunov: Data collection; Resources. Aysenil Belger: Data collection; Resources. Judy Ford: Data collection; Resources. Sarah McEwen: Data collection; Resources. Daniel Mathalon: Data collection; Resources. Bryon Mueller: Data collection; Resources. Godfrey Pearlson: Data collection; Resources. Steven G. Potkin: Data collection; Resources. Adrian Preda: Data collection; Resources. Jessica Turner: Data collection; Resources. Theodorus Van Erp: Data collection; Resources. Jing Sui: Writing – review & editing. Vince Calhoun: Conceptualization; Funding acquisition; Investigation; Methodology; Project administration; Resources; Writing – review & editing.

## Appendices

